# Non-destructive collection and metabarcoding of arthropod environmental DNA remained on a terrestrial plant

**DOI:** 10.1101/2022.09.27.509650

**Authors:** Kinuyo Yoneya, Masayuki Ushio, Takeshi Miki

## Abstract

For the conservation and community ecology of arthropods and pest controls on terrestrial plants, survey of arthropods is a crucial step. However, efficient surveys are hindered by challenges in collecting small arthropods, and identifying the species. Environmental DNA (eDNA)-based techniques, such as eDNA metabarcording, help overcome these difficulties in aquatic systems. To apply eDNA metabarcording to terrestrial arthropods, we developed a non-destructive eDNA collection method. In this method, termed “plant flow collection,” we spray distilled or tap water, or use rainfall, which eventually flows over the surface of the plant, and is collected in a container that is set at the plant base. DNA is extracted from the filtered water samples and a DNA barcode region, such as cytochrome c oxidase subunit I (COI) gene, of the extracted DNA is amplified and sequenced using a high-throughput sequencer. We identified more than 38 taxonomic groups of arthropods at the family level, of which 7 were visually observed or artificially introduced species, whereas the other 31 groups of arthropods, including 18 species, were not observed in the visual survey. Put together, these results show that the developed method is effective for detecting the presence of arthropods in plants.

## Introduction

Arthropod species in terrestrial ecosystems are diverse ^[1]^. The diversity of arthropods is influenced by the diversity of plants and their interactions with the plants that they consume and live on ^[2,3]^. The survey of various arthropods is a crucial step in studies of the conservation and community ecology of arthropods on plants and pest management in agriculture. Researchers have invested considerable efforts in identifying and exploring arthropods. Arthropods frequently have small bodies and occasionally exhibit limited morphological variation; thus, distinguishing the juvenile stages is difficult, which prevents rapid and accurate identification. Therefore, taxonomic expertise is necessary for morphology-based identification, but the number of amateurs and professional taxonomists is decreasing^[4]^. In addition, monitor them is time-consuming and costly^[5]^ because of their hiding behavior, concealed coloration, and relatively high mobility.

DNA metabarcoding, which combines universal primers and high-throughput sequencing, enables us to estimate how diverse species are, and is promising to overcome the difficulty in species identification. The cytochrome c oxidase I (COI) region of mitochondrial DNA has been frequently used for species identification of arthropods^[6,7]^. Environmental DNA (eDNA) metabarcoding has recently gained attention as a tool for efficient biodiversity monitoring, particularly in aquatic systems^[8]^. eDNA is genetic material originating from organisms, such as metabolic waste and tissues from the body surface^[9,10]^. It has been shown to be useful for investigating species that are difficult to find because of low population density or its behavior (e.g., nocturnal and quick-moving behavior)^[11,12]^. eDNA metabarcoding enables us to estimate the diversity of species in a certain water area^[13]^. Thus, eDNA metabarcoding could contribute to ecological research and biota monitoring over a wide area and in the long term^[10,13]^. eDNA metabarcoding has also been applied in terrestrial ecosystems to detect mammals living in forest ecosystems ^[14]^ and honeybees ^[15]^. However, eDNA metabarcoding has not yet been applied to organisms associated with terrestrial plants because a method to collect eDNA remaining on plant surfaces is not as well developed as that for aquatic eDNA. In several cases, the eDNA of herbivores remaining on plants has been successfully detected using destructive methods^[16–19]^. For example, ungulate browsing preference was investigated by analyzing animal eDNA in saliva that adhered to a bite site on twigs^[17,18]^.

A method to collect eDNA from plants, such as crops and endangered or protected plants, without damaging them would be preferred. Several studies have proposed potentially noninvasive methods for collecting eDNA from terrestrial plants. For example, to detect an invasive spotted lanternfly, *Lycorma delicatula*, eDNA was collected using a cotton roller and sprayed with water on a part of a plant^[20]^. However, eDNA metabarcoding of arthropod communities on the whole plant body has not been performed using non-destructive methods for both plants and arthropods.

In the present study, we investigated a non-destructive method for the eDNA metabarcoding of arthropod communities that feed and live on a plant. As in other studies, we focused on the utility of water as a collection medium for terrestrial eDNA ^[19,20]^. To this end, we sprayed distilled or tap water on the whole body of an eggplant and cabbage growing on a pot or on the ground in a field and collected the water flow at the base of plants, assuming that the water running through the whole plant body may contain arthropod DNA remaining on the plant. Because we aimed to disseminate this technique to detect pest insect attacks and apply it to large-scale biodistribution surveys (e.g., with citizen participants), we tested tap water for a simplified process in addition to distilled water for collecting eDNA. We also collected rainfall that had run through cabbage grown in a field as a potential alternative to manual water addition methods.

## Result & Discussion

Using eDNA metabarcoding, 38 taxonomic groups of arthropods, including those identified at least at the family level, were detected, belonging to 13 orders, 28 families, 30 genera, and 22 species (Table 1). Of these, seven were *A. gossypii, P. rapae, P. xylostella, M. persicae, B. brassicae*, a leaf miner fly (Agromyzidae), and *Sphaerophoria* sp, which were artificially introduced or visually observed on an experimental plant. They were hereafter termed “target species” in this study. An additional 31 arthropods were neither introduced nor observed by visual survey.

**Table 1.**
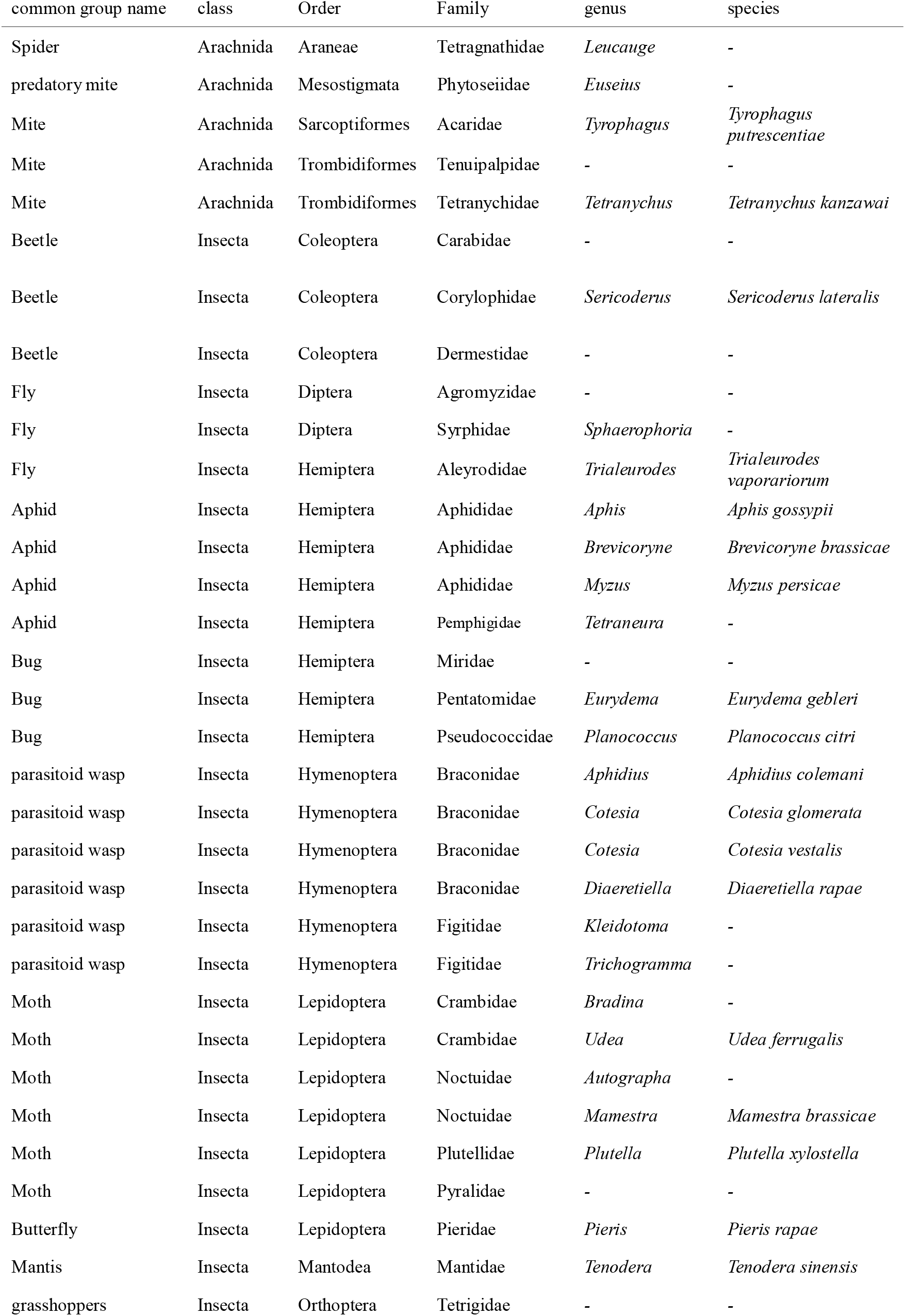
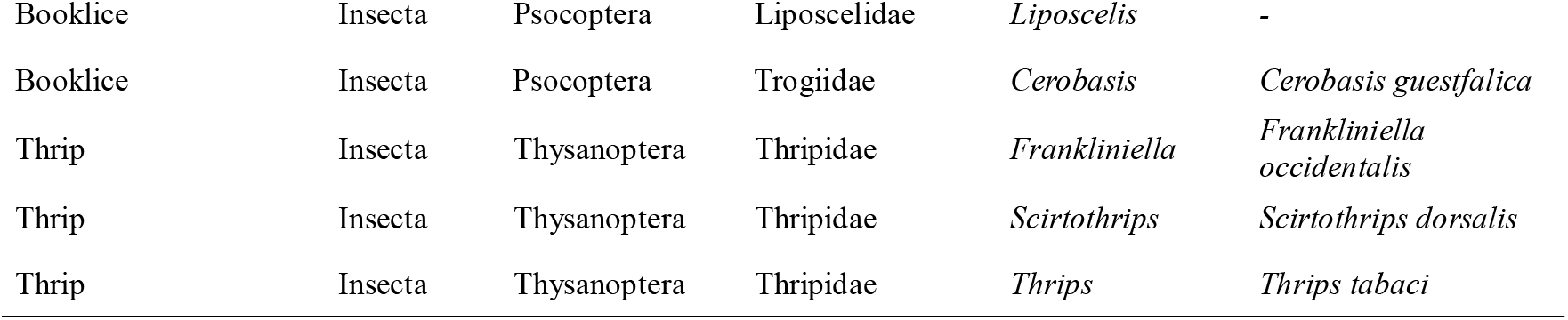
List of taxonomic information of arthropods identified in eDNA samples collected from plant surface using the “plant flow collection” method (see, fig. 2).

### eDNA collected from potted eggplant

Cotton aphids *A. gossypii* were detected in all eDNA samples collected from eggplants using tap water (3/3 samples, ID: S1-S3 in Table 2) and distilled water (2/2 samples, ID: S49, S50 in Table 2). Thus, both tap water and distilled water appeared to be effective media for eDNA collection (Table 2). It might be important to filter water samples as soon as possible after collecting eDNA using tap water because chloride in tap water can degrade the collected eDNA. Most eggplants were infested by leaf miners in our greenhouse this year, and many feeding tracks remained on the leaves. Thus, detection of leaf miner-fly Agromyzidae in all eDNA samples collected from eggplants in July (3/3 samples) is convincing (but not detected in October; 0/2 samples, Fig. S1). We also collected eDNA from eggplants with feeding tracks of leaf miners, but without herbivorous individuals, in August and September (see Methods and Table 3). Agromyzidae was detected in only one of four samples collected in August and not in September (0/4, Fig. S1). In July, when Agromyzidae was detected in all samples, Agromyzidae may have been engaged in feeding behavior on plants. On the other hand, in August and September, there were still traces of feeding damage, but Agromyzidae may have left most of the plants and eDNA may have already been degraded or washed away (e.g., see Valentin *et al*.^[27]^). We also detected several non-target species (i.e., individuals of several species could not be visually identified) in the samples collected from eggplants (Table S1). eDNA metabarcoding also detected major pest arthropods such as spider mites, *Tetranychus kanzawa* (10/12 samples), and a thrip, *Thrips tabaci* (7/12 samples, see ID: S1-3, S49,50 in Table S1). These herbivores are rarely controlled by tiny pests in agriculture because of their low visibility.

**Table 2.** The number of *Aphis gossypii* on a potted eggplant at the sampling day and total sequence reads detected from tap water or distilled water samples collected from plant surface by the “plant flow collection” method. The amount of water and sampling day was described in Table 3.

| Sample ID | No. of aphids | Water type*<sup>3</sup> | Sequence read |
| --- | --- | --- | --- |
| S1 | 39 | TW | 147 |
| S2 | 50 | TW | 54 |
| S3 | 157 | TW | 450 |
| S50 | Few C*<sup>2</sup> | DW | 22 |
| S49*<sup>1</sup> | Few C*<sup>2</sup> | DW | 314 |

\*<sup>1</sup>: Three egg plants were used for sampling. \*<sup>2</sup>: The number of aphids was not counted, but we chose plants with few colonies (C). \*<sup>3</sup>: Tap water (TW) or distilled water (DW) was used to collect eDNA. The water volumes and collection days are listed in Table 3.

**Table 3.**
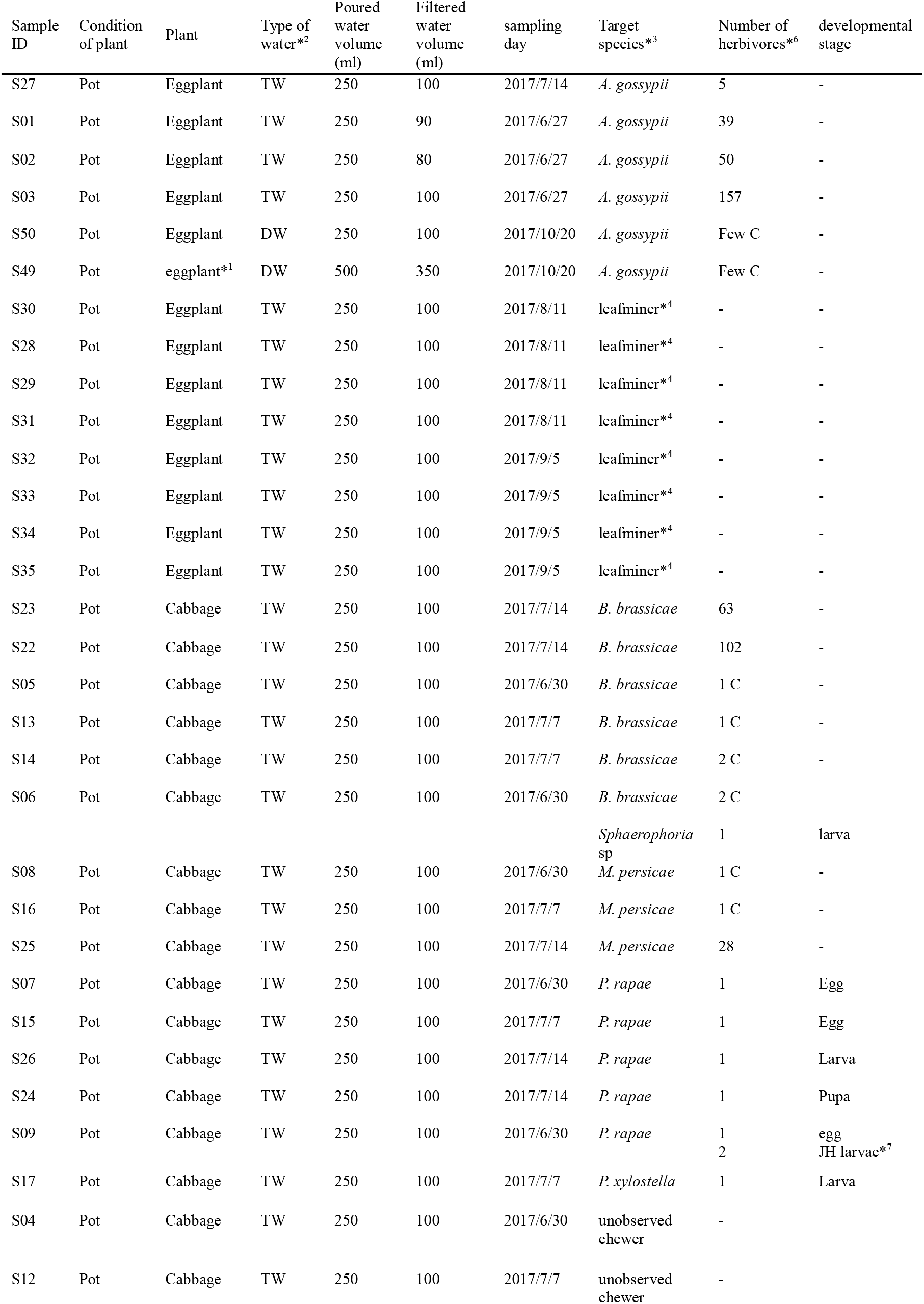
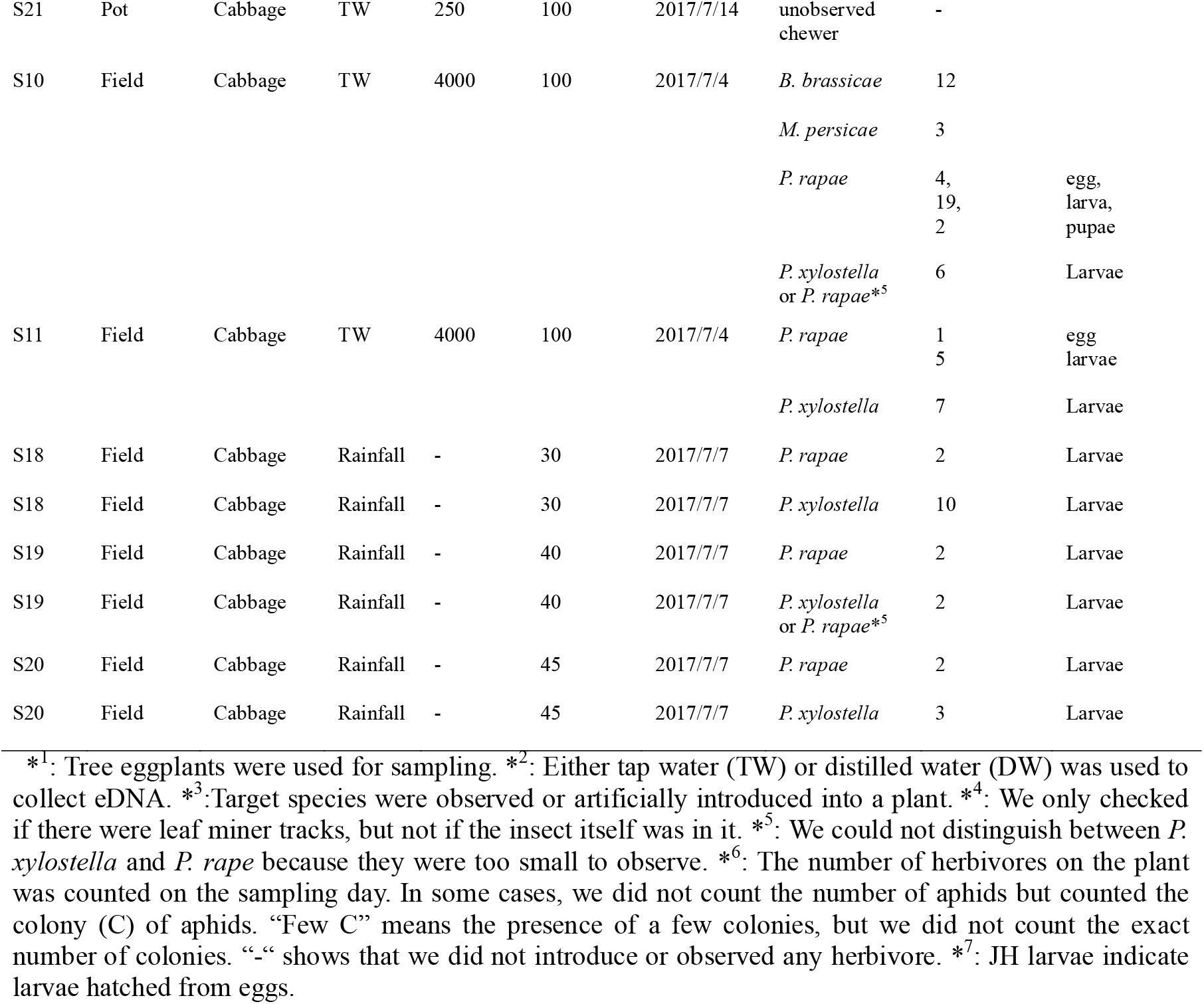
List of sample information.

### eDNA collected from potted cabbage

The eDNA of the target species *B. brassicae* and *M. persicae* was detected in water samples from potted cabbage plants (4/6 and 2/3 samples, respectively; Table 4). In many samples from which *B. brassicae* or *M. persicae* were detected, aphid parasitoids were also detected [*Diaeretiella rapae* (4/4 samples detected *B. brassicae*) and *Aphidius colemani* (2/4 samples detected *B. brassicae* and 1/2 samples detected *M. persicae*) (Table 4). In one of the eDNA samples collected from the cabbages, *Syrphidae sp*. was visually observed, in addition to *B. brassicae*, and its DNA was detected.

**Table 4.** The number of target species, *Brevicoryne brassicae, Myzus persicae*, and *Sphaerophoria* sp. observed on a potted cabbage at the sampling day and total sequence reads of these target species and the parasitoids, *Diaeretiella rapae* and *Aphidius colemani*, of these aphid species detected from samples collected by “plant flow collection”. Tap water was used to collect eDNA. The amount of water and sampling day was described in Table 3.

| Sample ID | Target species |  |  | Parasitoid of the target species |  |
| --- | --- | --- | --- | --- | --- |
|  | Species | The number | Sequence reads | Sequence reads of *D. rapae* | Sequence reads of *A. colemani* |
| S5 | *B. brassicae* | C* | 0 | 0 | 0 |
| S13 | *B. brassicae* | C* | 20598 | 4489 | 8 |
| S14 | *B. brassicae* | C* | 4 | 6 | 3 |
| S23 | *B. brassicae* | 63 | 379 | 950 | 0 |
| S22 | *B. brassicae* | 102 | 264 | 9445 | 0 |
| S6 | *B. brassicae* | C* | 0 | 0 | 0 |
|  | *Sphaerophoria* sp | 1 | 18 | - | - |
| S8 | *M. persicae* | C* | 41425 | 0 | 0 |
| S16 | *M. persicae* | C* | 0 | 0 | 0 |
| S24 | *M. persicae* | 28 | 18 | 0 | 6 |

\*The number of aphids was not counted, but we recorded that there were a few colonies (C).

*P. rapae* was detected in eDNA samples collected from potted cabbages with the three larvae, a larva or a pupa (S17, S26, and S25, respectively, Table 5), but not from a sample collected from a larva, an egg only, or an egg together with two larvae just hatched from the other eggs (S15, S7, and S9, respectively, Table 5). *P. xylostella* was also detected in eDNA samples (1/2 samples, Table 5). The detection probabilities of *P. rapae* eDNA seem to depend on the developmental stage, body size, and number of individuals on the plant.

**Table 5.** The developmental stage of target herbivorous species, *Pieris rapae* and *Plutera xylostella* observed on a potted cabbage at the sampling day and their sequence reads detected from 100 ml water samples collected from 250 ml tap water sprayed on plant surface by “plant flow collection”.

| Sample ID | Target species | The number of individuals | Developmental stage | Sequence reads |
| --- | --- | --- | --- | --- |
| S7 | *P. rapae* | 1 | Egg | 0 |
| S9 | *P. rapae* | 1 | Egg | 0 |
|  |  | 2 | & JH larvae* |  |
| S15 | *P. rapae* | 1 | Larva | 0 |
| S17 | *P. rapae* | 3 | Larva | 1324 |
|  | *P. xylostella* | 1 | Larva | 0 |
| S26 | *P. rapae* | 1 | Larva | 3 |
|  | *P. xylostella* | 1 | Larva | 6 |
| S25 | *P. rapae* | 1 | Pupa | 20 |

\* JH indicates larvae hatched from eggs.

Visually unobserved arthropods were also detected in eDNA samples from potted cabbage plants, such as tiny herbivore species, *T. kanzawai, T. tabaci*, and *Liposcelis* sp., grasshopper *Tetriginae* sp., leaf miner *Phytomyzinae* sp., and predator *Sphaerophoria* sp. (Table S1), in addition to the target species. Target species, *P. xylostella, P. rapae, M. persicae*, and *B. brassicae*, were also detected in some samples on which they were not visually observed. Similarly, several arthropods, such as *T. tabaci, B. brassicae*, and *Liposcelis*, were detected in eDNA samples collected from the surface of cabbage with chewing damage, on which no arthropods were visually observed (Table S2). In the three samples, chewers, which might have caused chewing damage, were detected in one sample (S12, in this case, *P. rapae*, Table S2) but not in the other two samples (S4 and S21, Table S2). This result suggests that to detect the eDNA of herbivores, it is not required that the herbivore exists on the plant when collecting the eDNA, but the longevity of eDNA on plant surfaces is limited.

### eDNA collected from field cabbage by using tap water and rainfall

Several arthropod species were detected in the eDNA samples collected from the surface of cabbage grown in the field by both rainfall and tap water (tap water: S10,11, rainfall: S18-20, fig. 1). Among these species, visually observed target species, such as *B. brassicae* and *P. rapae* (1/1 sample and 5/5 samples, respectively, fig. 1) were included. Because it was very difficult to distinguish young larvae of *P. rapa* and *P. xylostella* by eye, we could not calculate the exact detection probability of *P. xylostella* (approximately 3/5 samples, fig. 1). In addition, many unobserved arthropods were detected in eDNA samples. In some samples in which *B. brassicae* was detected, aphid parasitoids and *D. rapae* were also detected (2/3 samples were positive for *B. brassicae*, fig. 1). This result was similar to that obtained for potted cabbage (Table 4). Larval parasitoids of *P. rapae* and *P. xylostella, Cotesia glomerata* (3/5 samples detected *P. rapae*), and *C. vestalis* (1/3 samples detected *P. xylostella*) were also detected in eDNA samples, which were detected in lepidopteran larvae (fig. 1). In addition to the parasitoids, an egg parasitoid, *Trichogramma* sp., and a parasitoid of the leaf miner fly, *Kleidotoma* sp., were also detected (Table S1). Other visually unobserved arthropods were also detected [for example, tiny herbivores such as an onion thrip *T. tabaci*, a western flower thrip *Frankliniella occdentalis* (fig. 1), and a barklice *Sminthurinus* sp. (Table S1), lepidopterans *Manestra brassicae, Udea ferrugalis*, a stink bug *Eurydema gebleri*, and predators, a mantis *Tenodera sinensis* (fig. 1), and a spider *Leucauge* sp. (Table S1)]. *U. ferrugalis* may have been introduced by trees planted in the surrounding cabbage fields. However, we did not find any evidence that *U. ferrugalis* exists in Japan. DNA-based identification might be mistaken, and there is a possibility that this species is closely related to *U. testacea*, which feeds on cabbage.

**Figure 1.**
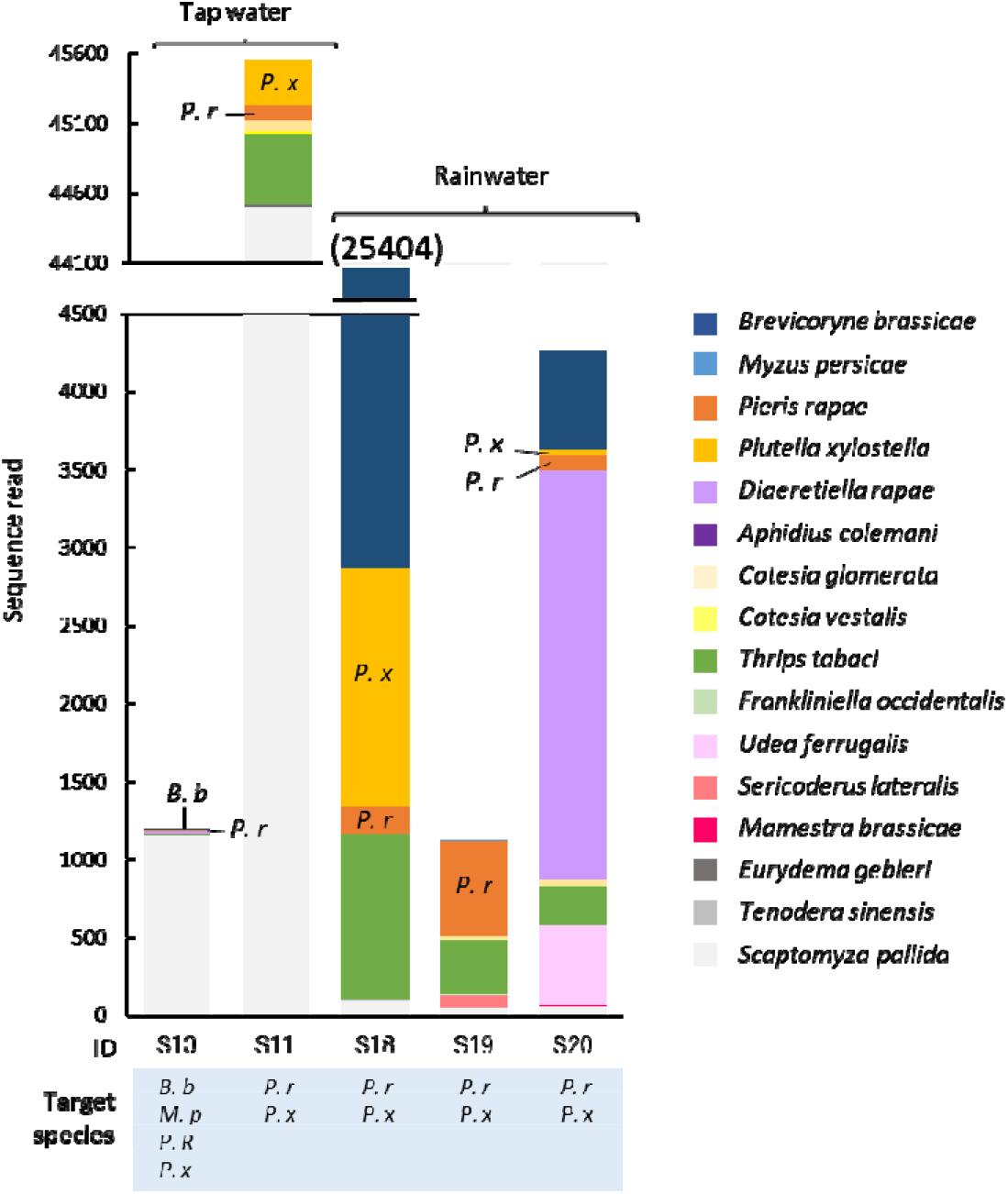
The sequence reads of species, which identified at species level by Claident or NCBI BLASTN programs, detected from eDNA samples collected from the surface of field cabbage by tap water (S10 and S11) or rainfall (S18-20). The other sequences identified at family or genus level are also described in Table S1. Label of under each bar is sample ID and the target species of each sample, *B. b*: *Brevicoryne brassicae, M. p*: *Myzus persicael, P. r*; *Pieris rapae, P*.*x*: *Plutella xylostella*. Target species were visually observed on a cabbage plant at the sampling day on July 4 (S10 and S11) and July 7, 2017 (S18-20). When target species were detected in a sample, the abbreviation of species name is described in a graph. The value in the parenthesis is the total sequence reads of S18. The number of target species observed on a sampling plant is described in Table 3.

### Detection of eDNA of visually unobserved species

There are two possible reasons for the detection of several arthropod species that were not visually observed on plants when the eDNA samples were collected. First, arthropods are simply too small and overlooked. The second reason is that they had existed on the plant previously and only eDNA remained on the plant (see Valentin *et al*.^[27]^ for the persistence of eDNA on plants). In future studies, it will be necessary to further investigate the “ecology” of eDNA (e.g., how it is released, moved, and persists on a plant surface), which is currently being extensively studied in fish eDNA studies^[28,29]^. In a previous study, eDNA of ungulate species was detected in 50% of ungulate species browsed twigs even after 12 weeks^[17]^. However, arthropods would release much less amount of their DNA on plants, so the eDNA of arthropods would become undetectable much faster than that of ungulate species.

## Conclusion

In this study, we developed a non-destructive method for collecting eDNA from whole plants by water spraying. It detected a wide range of arthropods, including not only conspicuous herbivores, but also parasitoids, predators, and tiny organisms, which are difficult to detect by eye, from whole plant bodies with a single sample collection. In some cases, small insects such as aphids or spider mites themselves may be included in the collected water, but including arthropod individuals themselves is not a matter for monitoring purposes. It is necessary to further investigate the differences in detection probabilities among different taxa of arthropods, number of individuals, and feeding behaviors. At the same time, our study clearly shows that, by applying our method of “plant flow collection” (i.e., collection of sprayed water or rainfall at the bottom of a plant, fig. 2), it is possible to detect small arthropods that are often overlooked, natural enemies of pest arthropods, such as parasitoids (i.e., *D. rapae* and *C. glomerata*) that are not easily observed visually, leaf miners that are hidden in leaves, and grasshoppers that are highly mobile. Therefore, our non-destructive method would be widely adaptable for surveying pests and natural enemies in pest management and monitoring arthropods in plant species.

**Figure 2.**
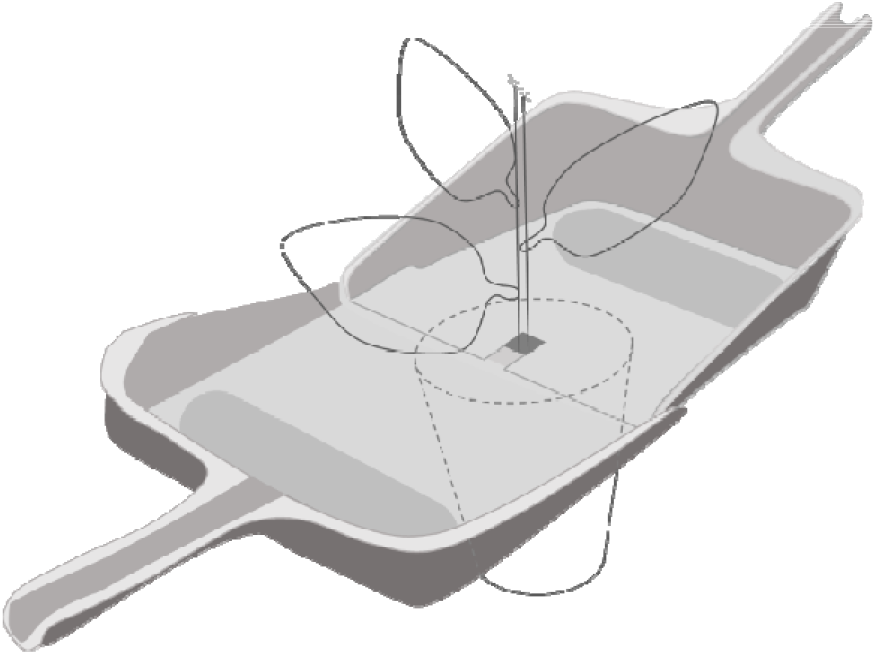
Pattern diagram of the “plant flow collection” method, which uses a set of two dustpans connected to each other. Water sprayed on and run through the whole body of plant was collected by this method. Dustpans were placed across the base of the plant stem on the soil.

## Methods

### Treatment for plants

The eggplants *Solanum melongena* L. and cabbage *Brassica oleracea* L. var. capitata L. were cultivated individually in pots (Φ 7 cm, 9 cm high) filled with culture soil from seeds in a glass house (25 °C ± 10 °C). We used potted eggplant and cabbage with approximately five leaves for the laboratory experiments. Cotton aphids, *Aphididae gossypii* Glover, from a colony maintained in our laboratory (Kindai University, Nara, Japan, 34.6694 N, 135.7381E) were placed on the leaves of an eggplant (5, 10, or 50 aphid individuals on each plant) in a transparent plastic box (25 cm × 30 cm × 28 cm) in a climate chamber (25 ± 3 °C, 16:8 h light: dark cycle) and kept for 7 d (the final number of aphids on a plant was 39, 50, and 157, respectively).

We selected potted cabbage with ca. 5 leaves in a glass house infested naturally by white butterfly *Pieris rapae* L, diamond back moth *Plutella xylostella* L., green peach aphid *Myzus persicae* Sulzer, or cabbage aphid *Brevicoryne brassicae* L., kept in a plastic case (27 × 15 × 13 cm), and collected eDNA from plant surfaces with these insects. We also used potted cabbage with only the feeding tracks of a leaf miner (N = 8) or chewers (N = 3) to test whether herbivores could be detected only from the feeding tracks. The sampling days are listed in Table 3.

Forty-eight cabbage plants with approximately five leaves per plant were transplanted in a field along with six lines. There was a 1 m interval between lines, and there were eight plants in each line with a 50 cm interval between plants in a common garden of the Faculty of Agriculture, Kindai University in Nara, Japan, in May 2017. Five randomly selected cabbages in the cabbage field were used for the experiments when they formed heads on July 4^th^ and 7^th^, 2017.

Seven arthropod species, *A. gossypii, P. rapae, P. xylostella, M. persicae, B. brassicae*, a leaf miner fly (Agromyzidae), and *Sphaerophoria* sp, which were artificially introduced or visually observed on an experimental plant, were hereafter termed “target species” in this study.

### Collection of eDNA

All sampling equipment was sterilized with a 10% bleach solution before use. A dustpan (1–4 cm depth × 25 cm wide × 20 cm with a 12 cm handle) with a cut (2 cm × 5 cm) opposite to the handle, where the stem of a cabbage or eggplant was in, and another uncut dustpan were connected to it (fig. 2) and covered with a polyvinylidene chloride lap to close the gap between the hole of the dustpan and the stem. Similarly, two dustpans covered with polyvinylidene chloride lap were inserted under cabbage planted in the field. Because we aimed to disseminate this technique to detect pest insect attacks and apply it to large-scale biodistribution surveys (e.g., with citizen participants), we tested tap water for a simplified process in addition to distilled water for collecting eDNA. Tap or distilled water was sprinkled throughout the plant body. Two-hundred fifty ml tap water or distilled water was used for each potted plant by a 250 mL wash bottle and 4 L tap water was sprayed using a 2 L watering can with a spray head or rainfall were used for field cabbage (Table 3). The sprayed water ran through the whole plant surface from the leaves to the stems, and water was gathered on the dustpan. Rainwater pooled over a day was collected one day after the dustpan was set under the cabbage. Details of the sample information, such as sampling day, amount of poured and filtered water, and the number and developmental stage of herbivores observed on a plant, are described in Table 3. We refer to this method as “plant flow collection”.

We collected 30–350 ml water (Table 3) pooled in dustpans by sacking with a plastic disposable syringe (SS-50LZ, Terumo Co.) and filtered with φ0.45 µm Sterivex™ filter cartridges (SVGV010RS, Merck Millipore, Darmstadt, Germany). Two milliliters of RNAlater solution were added to the filter cartridges and stored at −60 °C until further processing.

### DNA extraction

The RNAlater solution in each filter cartridge was removed by vacuum (EZ-Vac Vacuum Manifold) and then washed with 1 ml of MilliQ water. DNA was extracted from cartridge filters using a DNeasy® Blood & Tissue Kit (QIAGEN, Hilden, Germany) following the method described by Miya *et al*.^[21]^. Briefly, proteinase K solution (20 µL), PBS (220 µL), and buffer AL (200 µL) were mixed and 440 µL of the mixture was added to each filter cartridge. The materials on the cartridge filters were lysed by incubating on a rotary shaker (15 rpm) at 56 °C for 10 min. The mixture was transferred into a new 2 ml tube from the inlet of the filter cartridge by centrifugation (3,500 × g for 1 min). The collected DNA was purified using a DNeasy® Blood & Tissue Kit following the manufacturer’s protocol. After purification, the DNA was eluted using 100 µL of elution buffer provided with the kit. The eluted DNAs was stored at −20 °C until further processing.

### First and second PCR

Prior to library preparation, the workspaces and equipment were sterilized. Filtered pipette tips were used, and separation of pre- and post-PCR samples was carried out to safeguard against cross-contamination. Two negative controls (PCR-negative controls) were used to monitor contamination during the experiments. For the first PCR, 6 µL KAPA HiFi HotStart ReadyMix (KAPA Biosystems, Wilmington, WA, USA), 0.7 µL primer (5 µM of each primer; Forward, mlCOIIntF; Reverse, HCO2198^[22]^) with adaptor and six random bases (Table S3), 2.6 µL sterilized distilled H_2_O, and 2 µL DNA template. The thermal cycle profile after an initial 2 min denaturation at 95 °C was as follows (35 cycles): denaturation at 98 °C for 20 s, annealing at 52 °C for 30 s, and extension at 72 °C for 30 s, with a final extension at 72°C for 1 min. The first PCR product was purified using AMPure XP (PCR product: AMPure XP beads = 1:0.8; Beckman Coulter, Brea, California, USA). We performed a duplicate 1^st^ PCR. These products were pooled to reduce PCR dropouts, diluted 10-fold, and used as templates for the second PCR. 2^nd^ PCR was performed with 24 µl reaction volume containing 12 µl of 2 × KAPA HiFi HotStart ReadyMix, 1.4 µl of each primer (5 µM of each primer, Table S4), 7.2 µl of sterilized distilled H_2_O and 2.0 µl of template. Different combinations of forward and reverse indices were used for different templates (samples) for massive parallel sequencing with MiSeq (Table S5). The thermal cycle profile after an initial 2 min denaturation at 95 °C was as follows (12 cycles): denaturation at 98 °C for 20 s, annealing at 60 °C for 30 s, and extension at 72 °C for 1 min, with a final extension at 72 °C for 1 min.

Each 20 µL second PCR product was mixed together. The pooled library was purified using AMPure XP (PCR product: AMPure XP beads, 1:0.8). Appropriately 500 bp libraries were size-selected using 2% E-Gel Size Select (Thermo Fisher Scientific, Waltham, MA, USA). The double-stranded DNA concentration of the library was quantified using a Qubit dsDNA HS assay kit and Qubit fluorometer (Thermo Fisher Scientific, Waltham, MA, USA). The double-stranded DNA concentration of the library was adjusted to 4 nM using Milli-Q water, and the DNA was applied to the MiSeq platform (Illumina, San Diego, CA, USA). Sequencing was performed using a MiSeq Reagent Nano Kit v2 for 2 × 250 bp PE (Illumina, San Diego, CA, USA).

### Sequence data processing

All pipelines used for sequence read processing and taxonomic assignment followed Ushio^[23]^. Briefly, the raw MiSeq data were converted into FASTQ files using the bcl2fastq program provided by Illumina (bcl2fastq v2.18), and the FASTQ files were demultiplexed using Claident v0.2.2018.05.29 (http://www.claident.org^[24]^). Demultiplexed FASTQ files were analyzed using the Amplicon Sequence Variant (ASV) method implemented in DADA2 v1.7.0^[25]^. To filter the quality, forward and reverse sequences were trimmed to lengths of 240 and 200, respectively, based on the visual inspection of the Q-score distribution using the DADA2::filterAndTrim() function. The error rates were learned using the DADA2::learnErrors() function. Sequences were then dereplicated, error-corrected, and merged to produce an ASV-sample matrix. Taxonomic identification was performed for ASVs inferred using DADA2, based on the query-centric auto-k-nearest-neighbor (QCauto) method^[24]^. Taxonomic assignment was conducted with the lowest common ancestor algorithm^[26]^ using Claident v0.2.2018.05.29. The “semiall_genus” database (database containing all sequences in NCBI nt except sequences of vertebrates, *Caenorhabditis* and *Drosophila* from NCBI nucleotide sequences with genus or lower-level taxonomic information), which was prepared in Claident, was used for taxonomic assignment. The three classes, Insecta, Arachnida, and Collembola in Arthropods, were the focus of this study. The sequences were further identified by NCBI BLASTN programs if the species names of the target species (visually observed or artificially introduced on experimental plants) were not identified by Claident (Table S6).

## Acknowledgements

We thank Itsuki Jinno of Kindai University for his technical assistance. This work was supported by JSPS KAKENHI, a Grant-in-Aid for Young Scientists (B), Grant Number JP 17K15235 and by the Hakubi Project at Kyoto University (to MU).

## Author contributions

KY, MU conceived and designed the experiments. KY, TM performed the experiments. MU, TM analyzed the data. KY, MU, TM wrote the manuscript.

## Data availability statement

Submission No.= DRA014843

## Additional Information (including a Competing Interests Statement)

The authors declare no competing interests.

## References

[1] Stork, N. E., McBroom, J., Gely, C. & Hamilton, A. J. New approaches narrow global species estimates for beetles, insects, and terrestrial arthropods. Proc. Natl Acad. Sci. U. S. A. 112, 7519–7523 (2015).

[2] Ebeling, A. et al. Plant diversity effects on arthropods and arthropod-dependent ecosystem functions in a biodiversity experiment. Basic Appl. Ecol. 26, 50–63 (2018).

[3] Poelman, E. H., van Loon, J. J. A. & Dicke, M. Consequences of variation in plant defense for biodiversity at higher trophic levels. Trends Plant Sci. 13, 534–541 (2008).

[4] Hopkins, G. W. & Freckleton, R. P. Declines in the numbers of amateur and professional taxonomists: Implications for conservation. Anim. Conserv. 5, 245–249 (2002).

[5] Hegland, S. J., Dunne, J., Nielsen, A. & Memmott, J. How to monitor ecological communities cost-efficiently: The example of plant–pollinator networks. Biol. Conserv. 143, 2092–2101 (2010).

[6] Folmer, O., Black, M., Hoeh, W., Lutz, R. & Vrijenhoek, R. DNA primers for amplification of mitochondrial cytochrome c oxidase subunit I from diverse metazoan invertebrates. Mol. Mar. Biol. Biotechnol. 3, 294–299 (1994). PMID: 7881515.

[7] Brandon-Mong, G. J. et al. DNA metabarcoding of insects and allies: An evaluation of primers and pipelines. Bull. Entomol. Res. 105, 717–727 (2015).

[8] Taberlet, P., Bonin, A., Zinger, L. & Coissac, E. Environmental DNA: For Biodiversity Research and Monitoring (pnOxford Univ., 2018).

[9] Kelly, R. P., Port, J. A., Yamahara, K. M. & Crowder, L. B. Using environmental DNA to census marine fishes in a large mesocosm. PLOS ONE 9, e86175 (2014).

[10] Thomsen, P. F. & Willerslev, E. Environmental DNA - An emerging tool in conservation for monitoring past and present biodiversity. Biol. Conserv. 183, 4–18 (2015).

[11] Fukumoto, S., Ushimaru, A. & Minamoto, T. A basin-scale application of environmental DNA assessment for rare endemic species and closely related exotic species in rivers: A case study of giant salamanders in Japan. J. Appl. Ecol. 52, 358–365 (2015).

[12] Yonezawa, S. et al. Environmental DNA metabarcoding reveals the presence of a small, quick-moving, nocturnal water shrew in a forest stream. Conserv. Genet. 21, 1079–1084 (2020).

[13] Bohmann, K. et al. Environmental DNA for wildlife biology and biodiversity monitoring. Trends Ecol. Evol. 29, 358–367 (2014).

[14] Ushio, M. et al. Environmental DNA enables detection of terrestrial mammals from forest pond water. Mol. Ecol. Resour. 17, e63–e75 (2017).

[15] Utzeri, V. J. et al. Entomological signatures in honey: An environmental DNA metabarcoding approach can disclose information on plant-sucking insects in agricultural and forest landscapes. Sci. Rep. 8, 9996 (2018).

[16] Kudoh, A., Minamoto, T. & Yamamoto, S. Detection of herbivory: eDNA detection from feeding marks on leaves. Environ. DNA 2, 627–634 (2020).

[17] Nichols, R. V., Königsson, H., Danell, K. & Spong, G. Browsed twig environmental DNA: Diagnostic PCR to identify ungulate species. Mol. Ecol. Resour. 12, 983–989 (2012).

[18] Nichols, R. V., Cromsigt, J. P. G. M. & Spong, G. Using eDNA to experimentally test ungulate browsing preferences. Springerplus 4, 489 (2015).

[19] Valentin, R. E., Fonseca, D. M., Nielsen, A. L., Leskey, T. C. & Lockwood, J. L. Early detection of invasive exotic insect infestations using eDNA from crop surfaces. Front. Ecol. Environ. 16, 265–270 (2018).

[20] Valentin, R. E. et al. Moving eDNA surveys onto land: Strategies for active eDNA aggregation to detect invasive forest insects. Mol. Ecol. Resour. 20, 746–755 (2020).

[21] Miya, M. et al. Use of a filter cartridge for filtration of water samples and extraction of environmental DNA. J. Vis. Exp. (117), e54741–e54741 (2016).

[22] Leray, M. et al. A new versatile primer set targeting a short fragment of the mitochondrial COI region for metabarcoding metazoan diversity: Application for characterizing coral reef fish gut contents. Front. Zool. 10, 34 (2013).

[23] Ushio, M. Use of a Filter Cartridge Combined with Bead Beating Improves Detection of Microbial DNA from Water Samples 1–15, (2018).

[24] Tanabe, A. S. & Toju, H. Two new computational methods for universal DNA barcoding: A benchmark using barcode sequences of bacteria, Archaea, animals, fungi, and land plants. PLOS ONE 8, e76910 (2013).

[25] Callahan, B. J. et al. DADA2: High-resolution sample inference from Illumina amplicon data. Nat. Methods 13, 581–583 (2016).

[26] Huson, D. H., Auch, A. F., Qi, J. & Schuster, S. C. MEGAN analysis of metagenomic data. Genome Res. 17, 377–386 (2007).

[27] Valentin, R. E., Kyle, K. E., Allen, M. C., Welbourne, D. J. & Lockwood, J. L. The state, transport, and fate of aboveground terrestrial arthropod eDNA. Environ. DNA 3, 1081–1092 (2021).

[28] Murakami, H. et al. Dispersion and degradation of environmental DNA from caged fish in a marine environment. Fish. Sci. 85, 327–337 (2019).

[29] Jo, T. & Minamoto, T. Complex interactions between environmental DNA (eDNA) state and water chemistries on eDNA persistence suggested by meta-analyses. Mol. Ecol. Resour. 21, 1490–1503 (2021).

