## Supporting Information for "Non-destructive collection and metabarcoding of arthropod environmental DNA remained on a terrestrial plant"

**Title**

Contents:

Figure S1. The total sequence reads of leaf-miner flies, Agromyzidae, in eDNA samples.

Table S1. Sum of sequence reads of all the same lowest taxonomic groups at least family level, which were identified by Claident or NCBI BLASTN programs.

Table S2. The list of species detected from surface of cabbage having chewing damage, and their sequence reads.

Table S3. Primers of the first PCR

Table S4. Primer for 2^nd^ PCR

Table S5. List of indexes and reads of each sample by each ASV. The sequence and identified taxonomic information of each ASV is described in table S6.

Table S6. List of sequences and taxonomic informations of each ASV.

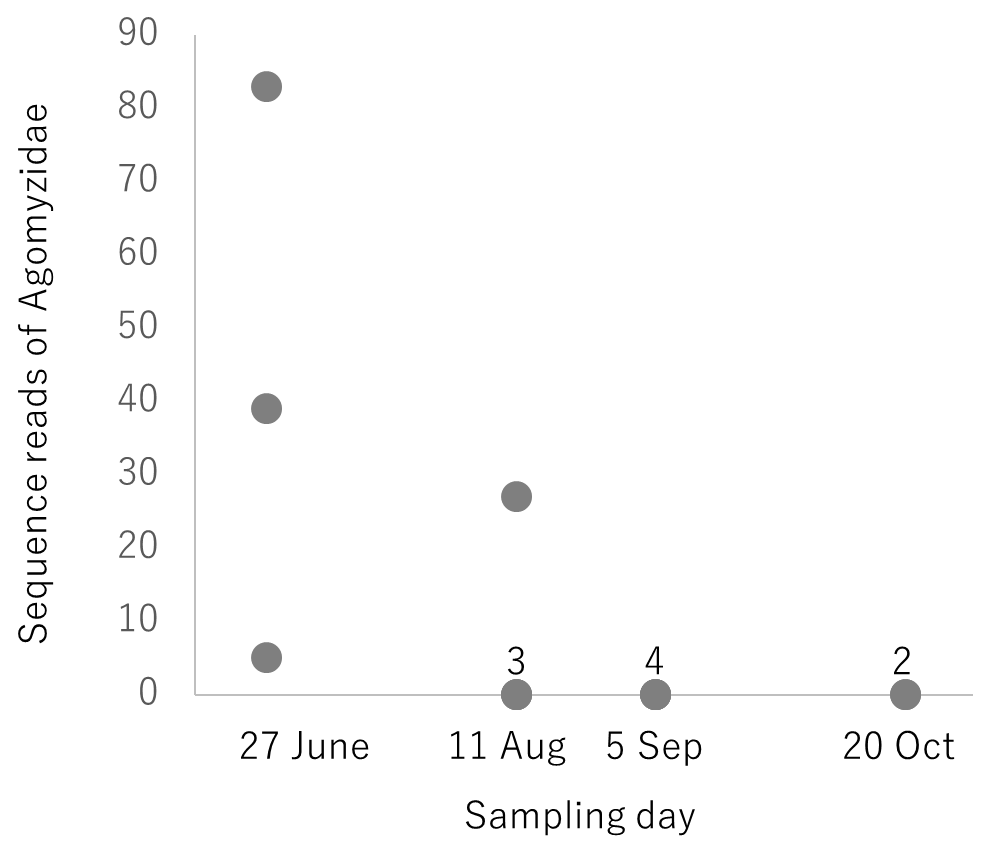

**Figure S1.** The total sequence reads of leaf-miner flies, Agromyzidae, in eDNA samples. The eDNA samples were collected from egg plants surface by the “plant flow collection” method at each sampling day in 2017. N = 3 (27 June, S1-3), 4 (11 August, S28-31), 4 (5 September, S32-35), 2 (20 October, S49,50). The numbers above plots are the number of samples showing the same 0 value.

Table S1. Sum of sequence reads of all the same lowest taxonomic groups at least family level, which were identified by Claident or NCBI BLASTN programs

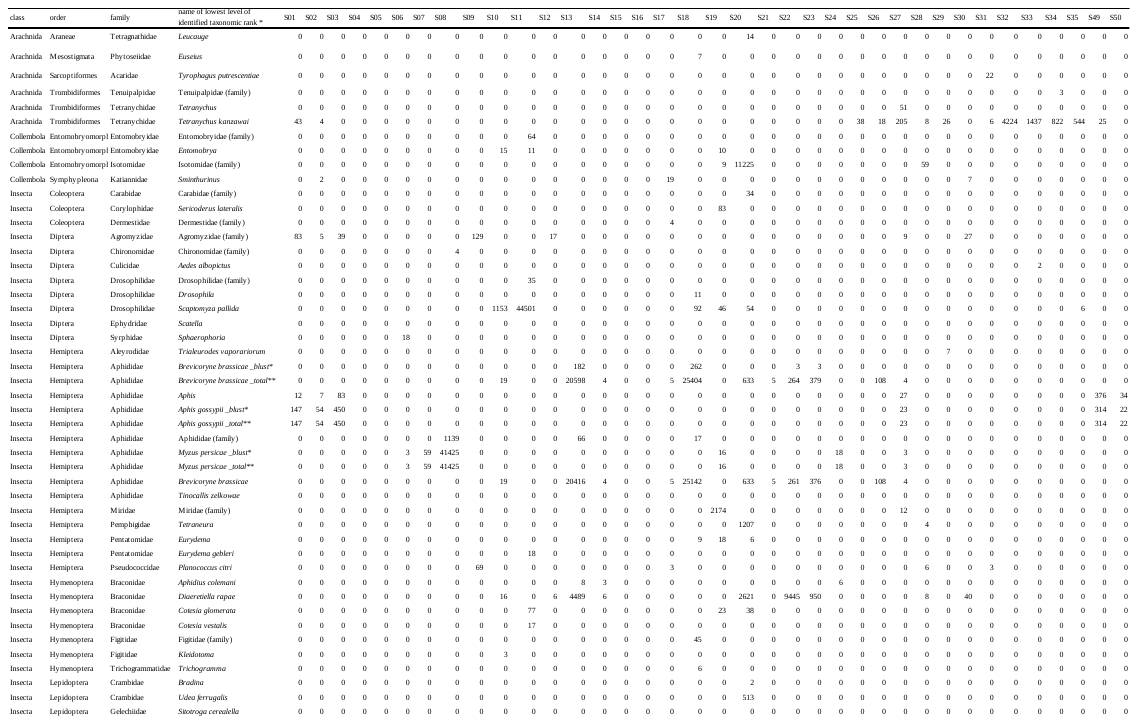

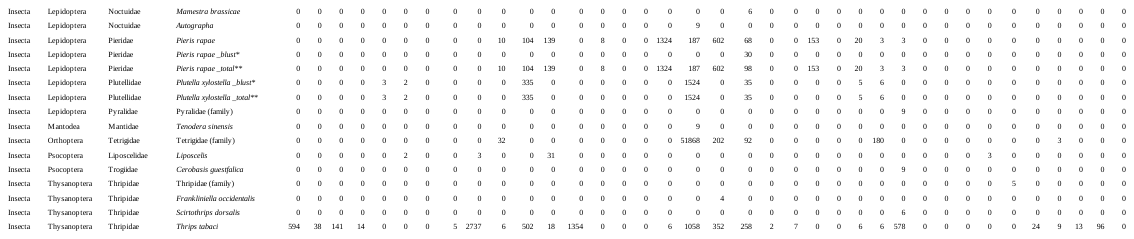

*_blust: the value is the total number of sequence reads of the species identified by NCBI BLASTN programs. **_total: the value is the total number of sequence reads of species identified by Claident and NCBI BLASTN programs.

Table S2. The list of species detected from surface of cabbage having chewing damage, and their sequence reads.

| Sample  ID | Species | sequence reads |
| --- | --- | --- |
| S4 | *Thrips tabaci* | 14 |
| S12 | *Pieris rapae* | 139 |
|  | *Diaeretiella rapae* | 6 |
|  | *Liposcelis* | 31 |
|  | *Thrips. Tabaci* | 18 |
|  | *Agromyzidae* | 17 |
| S21 | *Brevicoryne. brassicae* | 5 |
|  | *Thrips tabaci* | 2 |

Table S3. Primers of the first PCR

| **Name** | **1^st^ PCR primers (5´ -> 3´)*** |
| --- | --- |
| mlCOIintF | **ACACTCTTTCCCTACACGACGCTCTTCCGATCTNNNNNN**GGWACWGGWTGAACWGTWTAYCCYCC |
| HCO2198 | **GTGACTGGAGTTCAGACGTGTGCTCTTCCGATCTNNNNNN**TAAACTTCAGGGTGACCAAAAAATCA |

*Bold letters in the sequence indicate sequencing primer and six ambiguous sequences

Table S4. Primer for 2^nd^ PCR

|  | **2^nd^ PCR primers (5´ -> 3´)** |
| --- | --- |
| Forward | AATGATACGGCGACCACCGAGATCTACAC**XXXXXXXX**ACACTCTT TCCCTACACGACGCTCTTCCGATCT |
| Riverse | CAAGCAGAAGACGGCATACGAGAT**XXXXXXXX**GTGACTGGAGTT  CAGACGTGTGCTCTTCCGATCT |

*Bolded X indicates the index sequence. A list of index sequences is presented in Table S4.
